# Left Pulmonary Artery in 22q11.2 deletion syndrome. Echocardiographic evaluation in patients without cardiac defects and role of *Tbx1* in mice

**DOI:** 10.1101/517342

**Authors:** Gioia Mastromoro, Giulio Calcagni, Paolo Versacci, Carolina Putotto, Marcello Chinali, Caterina Lambiase, Marta Unolt, Elena Pelliccione, Silvia Anaclerio, Cinzia Caprio, Sara Cioffi, Marchesa Bilio, Anwar Baban, Fabrizio Drago, Maria Cristina Digilio, Bruno Marino, Antonio Baldini

**Affiliations:** Department of Pediatrics, Sapienza University of Rome, Rome; Department of Pediatric Cardiology and Cardiac Surgery, Bambino Gesù Pediatric Hospital and Research Institute, Rome; Institute of Genetics and Biophysics “Adriano Buzzati Traverso”, CNR, Naples; Department of Molecular Medicine and Medical Biotechnologies, University of Naples Federico II, Naples, Italy.

## Abstract

**Introduction and Hypothesis:** Patients with 22q11 deletion syndrome (22q11.2DS) present, in about 75% of cases, typical patterns of cardiac defects, with a particular involvement on the ventricular outflow tract and great arteries. However, in this genetic condition the dimensions of the pulmonary arteries (PAs) never were specifically evaluated.

We measured both PAs diameter in patients with 22q11.2DS without cardiac defects, comparing these data to a normal control group. Moreover, we measured the PAs diameter in *Tbx1* mutant mice. Finally, a cell fate mapping in *Tbx1* mutants was used to study the expression of this gene in the morphogenesis of PAs.

**Methods:** We evaluated 58 patients with 22q11.2DS without cardiac defects. The control group consisted of 54 healthy subjects, matched for age and sex. All cases underwent a complete transthoracic echocardiography. Moreover, we crossed *Tbx1*^*+/-*^ mice and harvested fetuses. We examined the cardiovascular phenotype of 8 wild type (WT), 37 heterozygous (*Tbx1*^*+/-*^) and 6 null fetuses (*Tbx1*^*-/-*^). Finally, we crossed *Tbx1*^*Cre/+*^mice with *R26R*^*mT-mG*^ Cre reporter mice to study *Tbx1* expression in the pulmonary arteries.

**Results:** The echocardiographic study showed that the mean of the LPA/RPA ratio in 22q11.2DS was smaller (0.80 ± 0.12) than in controls (0.97 ± 0.08; p < 0.0001).

Mouse studies resulted in similar data as the size of LPA and RPA was not significantly different in WT embryos, but in *Tbx1*^+/-^ and *Tbx1*^-/-^ embryos the LPA was significantly smaller than the RPA in both mutants (P=0.0016 and 0.0043, respectively). We found that *Tbx1* is expressed near the origin of the PAs and in their adventitia.

**Conclusions:** Children with 22q11.2DS without cardiac defects show smaller LPA compared with healthy subjects. Mouse studies suggest that this anomaly is due to haploinsufficiency of *Tbx1*. These data may be useful in the clinical management of children with 22q11.2DS and should guide further experimental studies as to the mechanisms underlying PAs development.

## INTRODUCTION

Patients with 22q11.2 deletion syndrome (22q11.2DS) present specific conotruncal defects (1) (2, 3) including tetralogy of Fallot with or without pulmonary atresia (4, 5), Truncus Arteriosus (6, 7), Interrupted Aortic Arch (8, 9), other aortic arch anomalies or minor congenital heart defects (10, 11), and ventricular septal defect (12).

In more than 90% of them a 3 Mb deletion was detected (13), spanning LCR22-A to LCR22-D that contains at least 30 genes including *TBX1* in the proximal region. This encodes a T-box transcription factor identified as the major player of this syndrome throughout both modeling mice (14-17) and mutational analysis in patients (18). In *Tbx1* mutant mice some cardiovascular anomalies similar to those found in 22q11.2DS patients have been described (15, 19). These observations can be explained by the fact that *Tbx1* is expressed in precursors of outflow tract cells and its loss of function reduces cell contribution to the outflow tract. (20–22).

Additional anomalies of pulmonary arteries (PAs) including diffuse hypoplasia, discontinuity, and crossing, were sporadically reported in 22q11.2DS patients (1, 3, 23-26) but not extensively studied. The aim of this study is to investigate the dimensions of both PAs in patients with this syndrome. In addition, we have analyzed the PAs diameters in *Tbx1* knockout mice and found that its haploinsufficiency is associated PAs asymmetry,indicating that this gene is the candidate for the PA phenotype reported here.

## MATERIALS AND METHODS

This is a prospective multicentric observational and experimental study conducted in three different Italian Centers: Department of Pediatrics, Sapienza University of Rome, Bambino Gesù Children’s Hospital and Research Institute, and Institute of Genetics and Biophysics of National Research Council, Naples.

Mouse studies were carried out at the Institute of Genetics and Biophysics under the auspices of the animal protocol 257/2015-PR (licensed to the AB lab) reviewed, according to Italian regulations, by the Italian Istituto Superiore di Sanità and approved by the Italian Ministero della Salute. The laboratory applies the “3Rs” principles to minimize the use of animals and to limit or eliminate suffering.

### Echocardiographic study

Data patients were ascertained from our hospital database for patients attending the Pediatric Cardiology Division of Sapienza University and Bambino Gesù Children Hospital. from October 2010 to April 2017. Informed consent was obtained from each patient (or legal guardians) and the study protocol conforms to the ethical guidelines of the 1975 Declaration of Helsinki. as reflected in a priori approval by the Institution’s Human Research Committee.

We included 58 pediatric and adult patients with 22q11.2DS without intracardiac malformations. Six of them presented isolated non-obstructive abnormalities of aortic arch or epiaortic vessels. We excluded from our cohort subjects with cardiac defects because of possible flow-related bias in our PAs measurements. The group of 58 cases consisted of 23 females (39.6%) and 35 males (60.4%), mean age of 12.8 ±10 years, with a mean body surface area (BSA) of 1.17±0.5; our control group consisted of 54 subjects (41% females) with a mean age of 10.8 ±10 years and a mean BSA of 1.18±0.5.

All patients underwent genetic counseling, and fluorescent in situ hybridization was performed to confirm the specific microdeletion. Echocardiographic measurements were compared with healthy subjects matched for age, sex, and BSA. All patients and healthy controls underwent a complete transthoracic echocardiographic examinations using GE Vivid E9 (Medical Systems, Oslo, Norway) with M6SD and 7S convex probe and Philips Ie33 Machine (Philips Medical Systems, Andover, MA) with X-5 and X-7 probes. M Mode, 2-Dimensional, and Doppler examinations were performed in all subjects. In particular, pulmonary branches were measured in parasternal short axis view during systole. Aortic arch anomalies were diagnosed in jugular view.

According to BSA, Z score values were reported for M Mode results and for 2D PA branches diameters. Images were digitally stored and measurement were made offline according to the American Society of Echocardiography guidelines by two independent readers for both centers (GM, PV, and GC, EP).

### Mouse studies

To perform phenotypic analyses, *Tbx1*^*+/-*^ mice (15) were intercrossed and pregnant females (3-6 months-old) were sacrificed using CO2 inhalation at plug day (E) 18.5, and fetuses harvested. Prior to observation. fetuses were washed in PBS and dissected under a Zeiss Stemi 2000-CS Stereo Microscope. Photographs were taken using a Z-stack software. In order to improve the view, we injected ink into the pulmonary trunk. Overall we have dissected and examined the cardiovascular phenotype of 8 wild type (WT), 37 heterozygous (*Tbx1*^+/-^), and 6 null fetuses (*Tbx1*^-/-^). To reveal *Tbx1*-expressing cells and their descendants. we crossed *Tbx1*^*Cre*/+^ mice (27) with *Rosa*^*mT*-*mG*^ mice. a Cre reporter (28). Hearts of E18.5 *Tbx1*^*Cre*/+^;*Rosa*^*mT*-*mG*^ embryos were dissected, photographed whole mount under Stemi 2000-CS Stereo Microscope with epifluorescence illumination, and then processed for cryosectioning. Sections were immunostained with an anti PECAM1 antibody (mouse monoclonal 2H8, Thermo Fisher MA3105, diluted 1:200) and/or an anti GFP antibody (Abcam ab13970, 1:800). Sections were photographed using a Leica fluorescence microscope. Digital images were mounted using Photoshop to generate the figures shown here.

## RESULTS

### 22q11.2DS patients have smaller LPAs than controls, independently from intracardiac anomalies

We identified 58 patients with isolated abnormalities of aortic arch or epiaortic vessels disease according to our criteria. Table 1 summarizes major clinical findings. All 22q11.2DS patients had variable expressivity and incomplete penetrance of dysmorphic features typical of the syndrome. A similar group of healthy volunteers was analyzed (Table 2). Using echocardiography, we have measured the diameter of PAs in a healthy subgroup and we found that LPAs were smaller than RPAs (9.4± 2.4 vs 10.0±2.8. P < 0.05). This finding was confirmed also in in 22q11.2DS cases, which exhibited LPAs measurements smaller than RPAs (8.2± 2.4 vs 9.4±2.4; P=0.016) and in Z score (−1.57± 1.2 vs −0.65± 1.0; P<0.001). No differences were found comparing diameters of RPAs in cases and controls. In contrast, LPA/RPA ratios showed a significant difference between the two groups: 0.80±0.09 in cases vs 0.95±0.11 in controls (P < 0.001) (Tabs. 1-2).

**Table 2.** Echocardiographic data of control patients.

| Case | Sex | Ao Arch<br>Anomalies | PAs<br>Anomalies | Age | Weight<br>(kg) | Height<br>(cm) | BSA | LPA d | z-score LPA | RPA d | z-score RPA | LPA/RPA |
| --- | --- | --- | --- | --- | --- | --- | --- | --- | --- | --- | --- | --- |
| 1 | M |  |  | 0 | 9 | 74 | 0.44 | 6.4 | 0.06 | 7.3 | 0.14 | 0.88 |
| 2 | M |  |  | 1 | 12 | 82 | 0.53 | 6 | -1.26 | 6.2 | -1.89 | 0.97 |
| 3 | F |  |  | 1 | 12.5 | 87 | 0.55 | 5.9 | -1.54 | 6 | -2.28 | 0.98 |
| 4 | F |  |  | 1 | 5.7 | 70 | 0.33 | 9 | 2.88 | 8 | 1.58 | 1.12 |
| 5 | F |  |  | 2 | 10.7 | 47 | 0.4 | 8 | 1.52 | 8.6 | 1.29 | 0.93 |
| 6 | M |  |  | 2 | 15 | 110 | 0.67 | 7.2 | -1.06 | 7.3 | -1.84 | 0.99 |
| 7 | M |  |  | 2 | 11.2 | 90 | 0.53 | 6 | -1.22 | 6.8 | -1.28 | 0.88 |
| 8 | M |  |  | 3 | 15.5 | 95 | 0.64 | 7.5 | -0.67 | 7.5 | -1.53 | 1 |
| 9 | F |  |  | 3 | 15.5 | 102 | 0.66 | 7.2 | -1.01 | 7.5 | -1.63 | 0.96 |
| 10 | F |  |  | 3 | 14 | 90 | 0.6 | 6.7 | -1.06 | 7.2 | -1.47 | 0.93 |
| 11 | M |  |  | 4 | 21 | 110 | 0.8 | 6.4 | -2.37 | 6.8 | -2.9 | 0.94 |
| 12 | M |  |  | 4 | 19 | 106 | 0.75 | 9.6 | 0.28 | 9.8 | -0.45 | 0.98 |
| 13 | F |  |  | 4 | 15 | 98 | 0.64 | 7.6 | -0.57 | 7.8 | -1.26 | 0.97 |
| 14 | F |  |  | 4 | 17 | 108 | 0.71 | 8.1 | -0.56 | 8.6 | -1.06 | 0.94 |
| 15 | M |  |  | 4 | 14 | 103 | 0.63 | 6.2 | -1.72 | 6.6 | -2.21 | 0.94 |
| 16 | M |  |  | 4 | 23 | 100 | 0.81 | 10.1 | 0.32 | 12.2 | 0.62 | 0.83 |
| 17 | M |  |  | 3 | 18 | 104 | 0.72 | 7.9 | -0.77 | 8.6 | -1.12 | 0.92 |
| 18 | F |  |  | 5 | 22 | 123 | 0.86 | 7.6 | -1.55 | 7.5 | -2.51 | 1.01 |
| 19 | M |  |  | 5 | 29 | 126 | 1.01 | 9.7 | -0.49 | 11.9 | -0.09 | 0.81 |
| 20 | F |  |  | 5 | 26 | 124 | 0.95 | 7.4 | -1.95 | 7.7 | -2.59 | 0.96 |
| 21 | F |  |  | 5 | 20 | 117 | 0.8 | 8.5 | -0.67 | 7.5 | -2.3 | 1.13 |
| 22 | M |  |  | 7 | 27 | 121 | 0.96 | 8.5 | -1.1 | 10 | -0.79 | 0.85 |
| 23 | M |  |  | 8 | 27 | 138 | 1.01 | 8.4 | -1.4 | 8.5 | -2.2 | 0.99 |
| 24 | F |  |  | 8 | 25.5 | 129 | 0.95 | 8.4 | -1.21 | 7.9 | -2.45 | 1.06 |
| 25 | F |  |  | 9 | 32 | 128 | 1.07 | 7.4 | -2.23 | 7.5 | -3.02 | 0.99 |
| 26 | M |  |  | 9 | 40 | 139 | 1.25 | 11 | -0.11 | 11 | -0.96 | 1 |
| 27 | M |  |  | 9 | 30.8 | 138 | 1.08 | 8.5 | -1.5 | 9.3 | -1.7 | 0.91 |
| 28 | F |  |  | 8 | 35 | 142 | 1.17 | 10.4 | -0.35 | 13.2 | 0.26 | 0.79 |
| 29 | F |  |  | 10 | 45 | 150 | 1.37 | 9.9 | -0.86 | 9.9 | -1.76 | 1 |
| 30 | M |  |  | 11 | 66 | 162 | 1.74 | 12 | -0.15 | 13.4 | -0.81 | 0.89 |
| 31 | M |  |  | 11 | 42 | 158 | 1.35 | 10 | -0.78 | 11 | -1.09 | 0.91 |
| 32 | M |  |  | 11 | 42 | 140 | 1.28 | 12 | 0.37 | 13 | 0.01 | 0.92 |

**Table 1.**
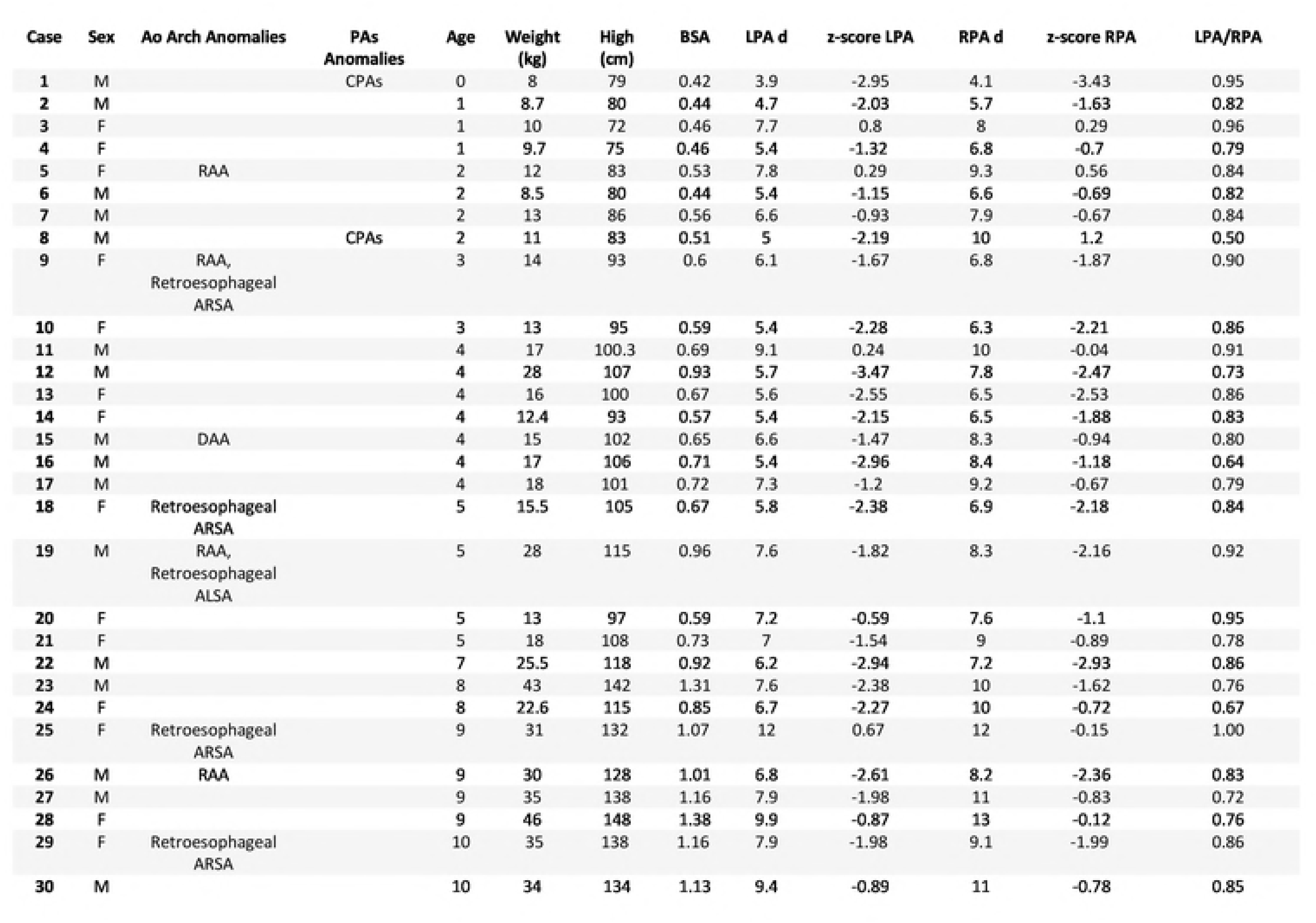
Echocardiographic data of 22q11.2DS patients without intracardiac defects.

### Tbx1 haploinsufficiency is associated with smaller LPAs in mice

*TBX1* is the candidate gene for many of the clinical and developmental features of 22q11.2DS patients including aortic arch anomalies and intracardiac anomalies. However, to our knowledge, anomalies of PAs in mouse mutants have not been reported to date. To understand whether loss of *Tbx1* may be a candidate also for the observed size asymmetry of the PAs, we measured them in *Tbx1*^+/+^, *Tbx1*^+/-^, and *Tbx1*^-/-^ E18.5 fetuses in a homogeneous congenic background C57Bl6/N. Results are plotted in Fig. 1. In WT (*Tbx1*^+/+^) fetuses, we found no significant difference (Mann-Whitney test) in the diameters of LPAs and RPAs (ratio LPA/RPA=0.92. n=8). However, in *Tbx1*^+/-^ fetuses, the LPAs were significantly smaller than the RPAs (P=0.0016, ratio LPA/RPA=0.79, n=37). Similarly, the *Tbx1*^-/-^ fetuses also had significantly different PAs (P=0.004, ratio LPA/RPA=0.63, n=6). All WT fetuses had normal arch and epiaortic vessels. Of the 37 heterozygous animals analyzed, 14 had aberrant origin of the right subclavian artery (37.8%) of which, 2 had high aortic arch, 3 interrupted aortic arch type B, and 1 right aortic arch. All 6 *Tbx1*^-/-^ fetuses had truncus arteriosus, as previously described (15). In all *Tbx1*^-/-^ fetuses the pulmonary arteries rose separately from the posterior wall of the arterial trunk proximal to the branches of aortic arch (Truncus Arteriosus - type II of Collett and Edwards or type A2 of Van Praagh).

**Figure 1.**
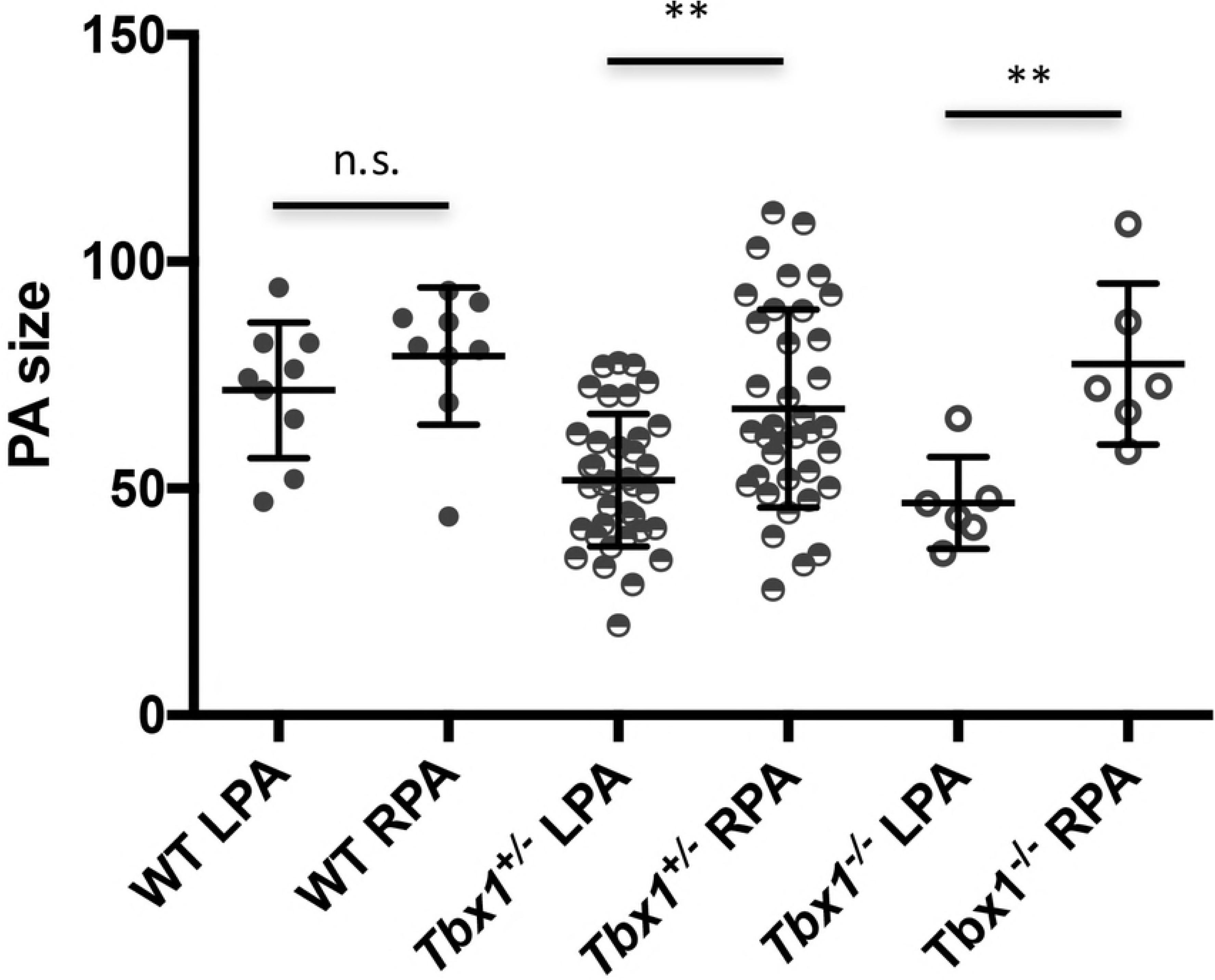
Distribution of pulmonary arteries measurements in WT, *Tbx1*^+/-^, and *Tbx1*^-/-^ fetuses at E18.5.

### Tbx1-expressing cells contribute to structural components of the pulmonary arteries

To provide insights as to how *Tbx1* may affect the development of the PAs, we looked into the expression of the gene. To do this. we used genetic marking of *Tbx1*-expressing cells and their descendants in *Tbx1*^*cre*/+^; *Rosa*^*mT*/mG^ embryos in which these cells are marked by membrane-bound green fluorescent protein (GFP). At E10.5. the PAs connect the aortic sac (through the proximal end of the 6th pharyngeal arch arteries) to the lung buds. GFP+ cells were observed in the mesoderm adjacent to the arteries, in the adjacent dorsal pericardial wall, and in the inner, endothelial layer of the arteries (Fig. 2). In the mature PAs at E18.5, the distribution of GFP+ cells were observed in the endothelial layer and in the outer mesenchymal tissue adjacent to the arteries. We did not observe contribution of GFP+ cells in the smooth muscle layer of the arteries, in contrast to the pulmonary trunk.

**Figure 2.**
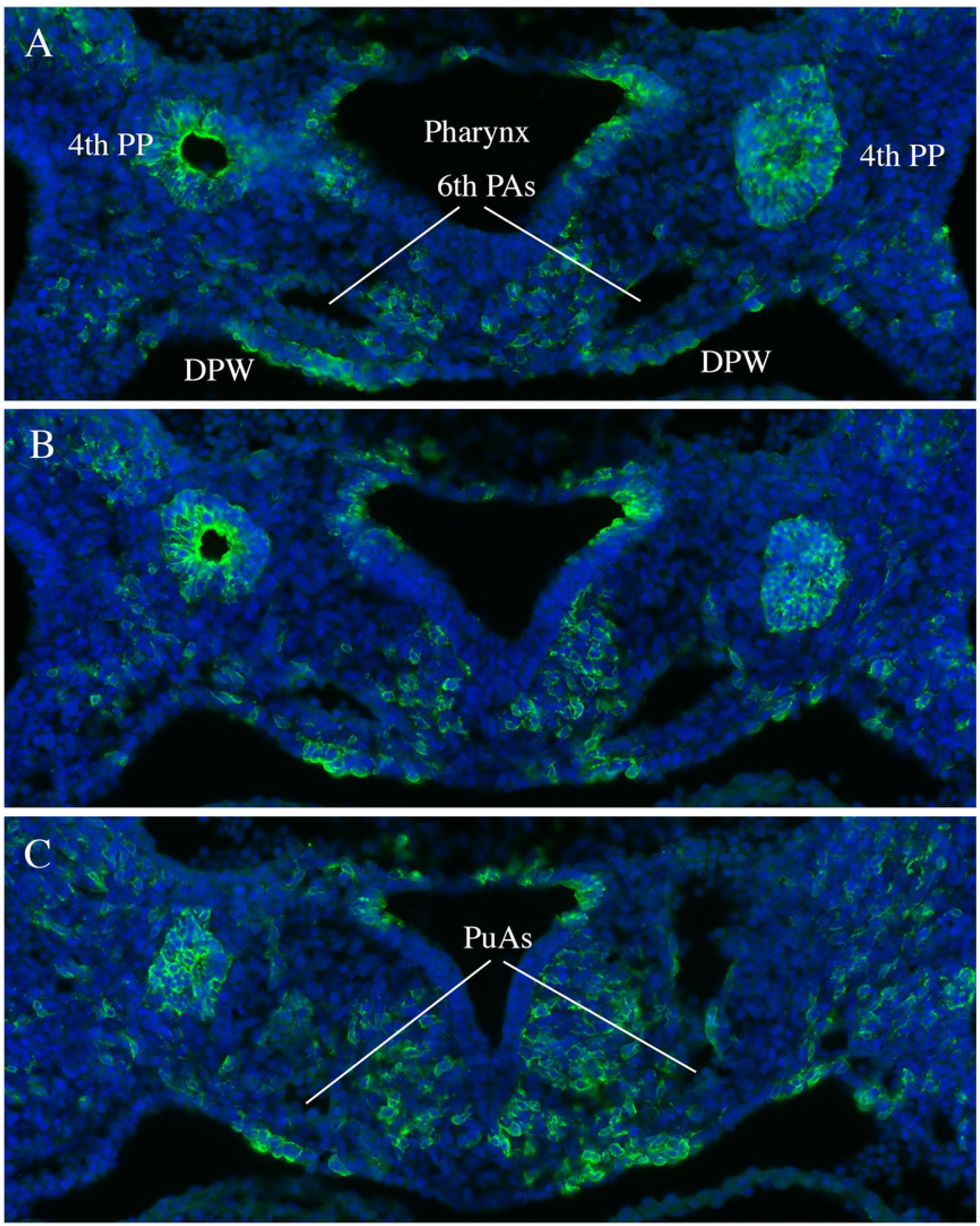
Transverse section of a E10.5 *Tbx1*^*cre*/+^; *Rosa*^*mT*/mG^ embryos immunostained with an anti GFP antibody (green). GFP positivity indicate cells that have expressed Cre recombinase. A, B, and C refer to 3 adjacent sections (cranial -> caudal) that span the junction between the 6th pharyngeal arch arteries (PAs) and the putative pulmonary arteries (PuA). PP: pharyngeal pouches; DPW: dorsal pericardial wall.

## DISCUSSION

The junction between LPA and DA is a crucial segment for the cardiovascular development and it is frequently affected in patients with conotruncal anomalies (29 - 32). Also in healthy people a smaller diameters of the LPA was reported in comparison with the RPA. Moreover, malformations of the pulmonary arteries, in particular of the left, including stenosis, diffuse hypoplasia, discontinuity or crossing, are not unusual in children with 22q11.2DS with or without conotruncal defects (33, 34, 3, 35, 36, 25, 24, 37). The detailed morphogenesis of the pulmonary arteries is not definitively ascertained. However, recent studies on mouse and human embryos contribute to better clarify this difficult topic (38, 39). According to recent data, while on the right side the VI pharyngeal arch artery disappears, on the left side it is formed by a ventral bud from the aortic sac and by a dorsal bud from the dorsal aorta. This ventral bud, with the contribution from the post-branchial pulmonary plexus, forms the LPA. The dorsal bud on the left side of the VI aortic arch forms the ductus arteriosus (DA), which is in continuity with the LPA. Our echocardiographic studies show that even in the absence of conotruncal defects, patients with 22q11.2DS have a smaller LPA compared to healthy subjects.

Our data demonstrate that the LPA is smaller than the RPA in *Tbx1*^+/-^ fetuses but not in WT fetuses, indicating that *Tbx1* haploinsufficiency affects significantly the LPA size. Expression data indicate that structural components of the PAs (endothelium and adventitia) derive from *Tbx1*-expressing cells. This is also true for the DA, thus suggesting that Tbx1 is involved in the development or growth of this cardiovascular segment. Mouse data are in agreement with echocardiographic measurements on patients with 22q11.2DS.

It is of interest to note that in mice *Tbx1* haploinsufficiency affects the IV but not the VI aortic arch artery development (19.40), while in *Tbx1*^-/-^ embryos the VI does not develop (19). The phenotype that we have described here suggest that a) the absence of the VI aortic arches does not have a dramatic impact on PAs development, and b) the reduced size of the LPA is probably not secondary to abnormalities of the VI aortic arch. The finding that Tbx1-expressing cells contribute to structural component of the PAs provides a support for a direct, though limited role of Tbx1 in determining the size of the PAs.

In the past, stenosis, diffuse hypoplasia or atresia of the proximal LPA was mainly ascribed to the extension of the ductal tissue into the LPA lumen. This pathogenetic mechanism known as “coarctation of the LPA” maintains its validity. However. our data suggest that molecular causes may influence the morphogenesis of this peculiar cardiovascular region, in particular the effect of *Tbx1* in this region may influence the morphology and dimensions of the LPA and its loss of function may cause some of its specific defects.

The reduced dimensions of LPA observed in our patients could be considered a subclinical sign associated with 22q11.2DS. We suggest that in subjects with 22q11.2DS the junction between the DA and the LPA may be at risk of additional anomalies and deserves specific diagnostic investigation.

## ACKNOWLEDGMENTS

This work was funded in part by grants from the Fondation Leducq (grant 15CVD01) and Telethon Foundation (grant GGP14211) to A. Baldini.

---

**Figure 3.**
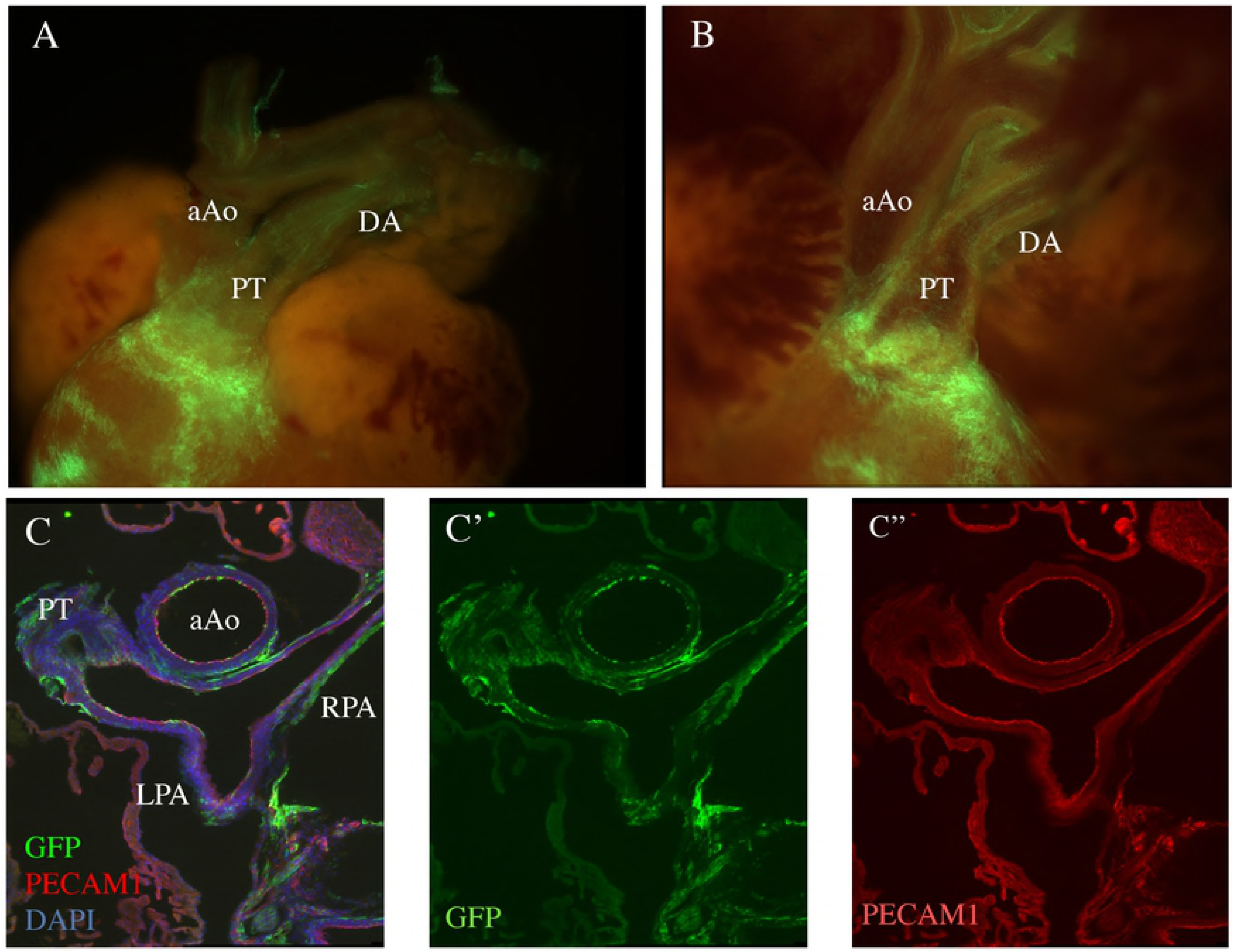
A.B: Whole mount fluorescent photographs of the outflow region of a E18.5 *Tbx1*^*cre*/+^; *Rosa*^*mT*/mG^ fetus. A: external appearance. B: internal optical plane. Note the heavy contribution of GFP+ cells to the pulmonary myocardium, pulmonary trunk (PT) and pulmonary valves, but superficial contribution (endothelial and adventitial) to other great vessels, including the ductus arteriosus (DA). which appears to have a more dense endothelial contribution. aAo: ascending aorta. C: immunofluorescence of a transverse section of a E18.5 *Tbx1*^*cre*/+^; *Rosa*^*mT*/mG^. Anti GFP staining is shown in green, anti PECAM1 staining (endothelial-specific) is shown in red. DAPI staining (cell nuclei) is shown in blu. C’-C’’: green and red channels are shown separately. aAo: ascending aorta; PT: pulmonary trunk and pulmonary leaflets; LPA, RPA: left and right pulmonary arteries.

